# The Phonetics-Phonology Relationship in the Neurobiology of Language

**DOI:** 10.1101/204156

**Authors:** Mirko Grimaldi

## Abstract

In this work, I address the connection of phonetic structure with phonological representations. This classical issue is discussed in the light of recent neurophysiological data which – thanks to direct measurements of temporal and spatial brain activation – provide new avenues to investigate the biological substrate of human language. After describing principal techniques and methods, I critically discuss magnetoencephalographic and electroencephalographic findings of speech processing based on event-related potentials and event-related oscillatory rhythms. The available data do not permit us to clearly disambiguate between neural evidence suggesting pure acoustic patterns and those indicating abstract phonological features. Starting from this evidence, which only at the surface represents a limit, I develop a preliminary proposal where discretization and phonological abstraction are the result of a continuous process that converts spectro-temporal (acoustic) states into neurophysiological states such that some properties of the former undergo changes interacting with the latter until a new equilibrium is reached. I assume that – at the end of the process – phonological segments (and the related categorical processes) take the form of continuous neural states represented by nested cortical oscillatory rhythms spatially distributed in the auditory cortex. Within this perspective, distinctive features (i.e., the relevant representational linguistic primitives) are represented by both spatially local and distributed neural selectivity. I suggest that this hypothesis is suitable to explain hierarchical layout of auditory cortex highly specialized in analyzing different aspects of the speech signal and to explain learning and memory processes during the acquisition of phonological systems.

## 1. Introduction

### 1.1 The linguistic perspective

In the beginning, phonetics and phonology did not constitute independent disciplines and their underlying notions were mutually interchangeable. With the emergence of the synchronic approach, phonetics and phonology were progressively separated: so, the assumption – immanent in all alphabetic writing systems – that the sounds of speech (*phones*) might be analyzable into separate units (*phonemes*) has taken root within contemporary linguistics (see Durand & Laks 2002; van der Hulst 2013 for a discussion). Both phonetics and phonology have the aim of describing and explaining sound patterns of human languages, but the way in which the two disciplines addressed this issue has been generally counterposed. So, the relationship between phonetics and phonology has been consistently gotten mixed up by puzzled.

Jakobson, Fant & Halle (1952)’s work prompted, in same way, the connection of phonetics substance with phonological mental representations. The crucial idea was that distinctive features (i.e., the abstract link between articulatory plans and acoustic outputs) are universal, binary, and must have correlates in terms of both articulation and audition: «The distinctive features would be more than a universal schema for classifying phonemes in all their diversity across languages; the features would be ‘real’ in the sense of being universal neural mechanisms for producing and for perceiving sounds of speech» (Jakobson & Waugh, 1979: 123). From this perspective, the relevant representational linguistic primitives are not single segments, but rather smaller segments are composed of: i.e., distinctive features. Accordingly, the work of the phonologist had to be interdisciplinary, as s/he needed to treating and interpreting data from language acquisition or loss, experimental phonetics, and psycholinguistics. However, the Jakobson’s approach remained programmatic and phonetics and phonology followed their own ways.

The *analysis by synthesis* framework offered another chance to reconcile the two levels of analysis according to Jacobson’s ideas (Stevens & Halle 1967; Stevens, 2002). The analysis by synthesis theory assumes top-down processes in which potential signal patterns are internally generated (synthesized) and compared to the incoming signal which is continuous and does not present markers for phonemes boundaries. Thus, perceptual analysis crucially contains a step of synthetically generated candidate representations (a form of hypothesis-and-test model). The model proceeds from the assumption that cues from the input signal trigger guesses about “landmarks” that serve to identify phoneme boundaries: consequently, the internal synthesis of potential phonemes is compared to the input sequence. Thus, landmarks are intrinsic to the signal and provide evidence for different kinds of segments (vowels, glides, and consonants): e.g., a peak in low-frequency amplitude for a vowel, a minimum in low-frequency amplitude, without acoustic discontinuities, for a glide, and two acoustic discontinuities for a consonant, one of which occurs at the consonant closure and one at the consonant release (Stevens, 2002: 1873; see also Poeppel et al., 2008).

For example, vowels may be classified on the basis of the first two formant peaks (F1, F2) on the spectral envelopes (Peterson & Barney 1952). The F1 is inversely correlated with articulatory tongue height, while the F2 (but also F3) reflects the place of articulation (PoA) along the horizontal (front-back and unrounded-rounded) dimension. The major features for vowels are the features specifying the position of the tongue body and lip rounding: the features [±high], [±low], [±back] and [±round] (as showed in Figure 1). In consonants, beyond formants, additional physical parameters are essential for discriminative performance: e.g., formant transitions, energy bursts, and the vibrations of the vocal chords occurring before and during the consonant burst.

**Fig. 1:**
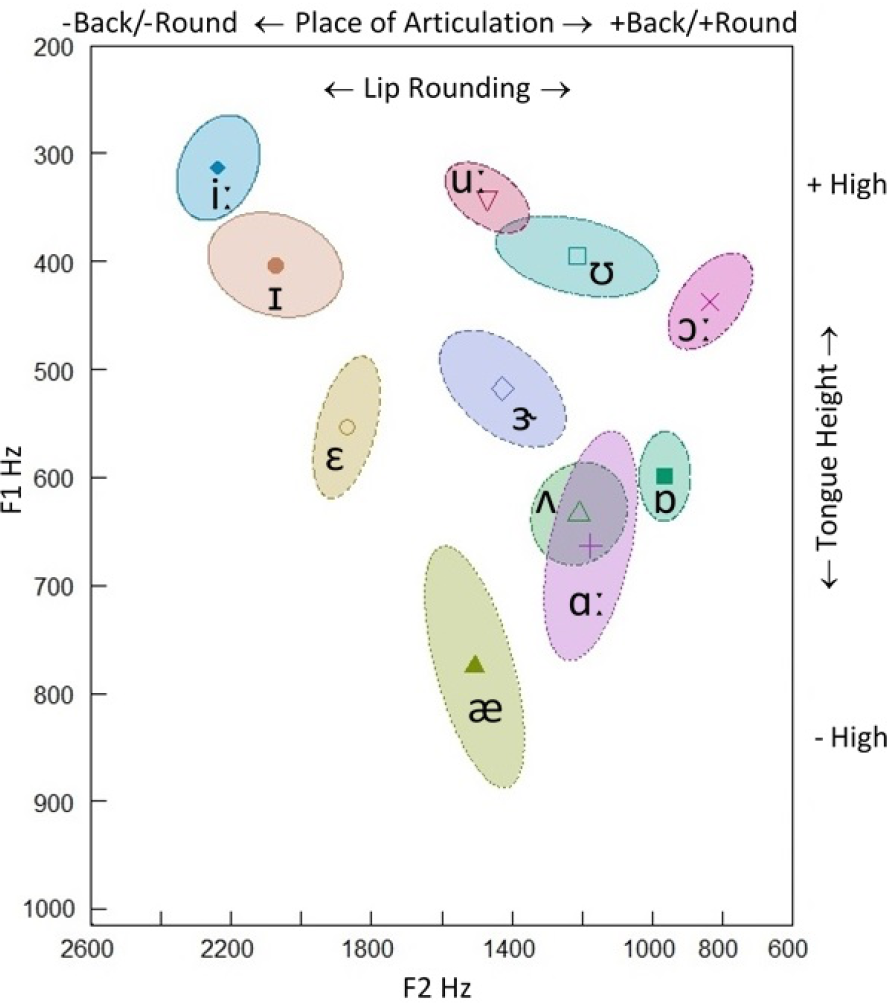
F1-F2 Hz scatterplot of the stressed British English vowels produced by a native 50-year-old male speaker (recorded at CRIL). 68.27% confidence ellipse corresponding to ±1 standard deviation from the bivariate mean (the symbol within the ellipse indicates the mean formant value). F1 is inversely correlated with articulatory tongue height (+high/−high), while F2 reflects place of articulation in the horizontal (−back/+back and −round/+round) dimension.

The publication of Chomsky & Halle (1968)’s volume sanctioned the separation of the two disciplines. This work emphasized the phonological component, that is, the system of rules, based on distinctive features, that applies to a surface structure and assigns to it a certain phonetic representation drawn from the universal class provided by linguistic universals. The phonological component is abstract (collocated at the level of the mental representation) as compared with phonetic representations, although both are given in terms of phonetic features. Therefore, the phonological principles that speakers acquire and control determine the phonetic shape of phonemes, words, and sentences. Words are represented as a series of phonemes each of which is a bundle of distinctive features that indicate the acoustic-articulatory configuration underlying phonological segments. In principle, the framework initially developed did not consider that the subject matter of phonology should be empirically verified since distinctive features (and their implications) were taken for granted (but see Halle 2002). This mainstream approach survived notwithstanding, for example, Halle (1983: 94-95) clearly suggested that distinctive feature representation comes both from articulatory and from acoustic/auditory aspects of speech sounds and that they are instantiated in the brain: «[…] we spoke not of ‘articulatory features’ or of ‘acoustic features,’ but of ‘articulatory’ and/or ‘acoustic correlates’ of particular distinctive features. […] On this view, the distinctive features correspond to controls in the central nervous system which are connected in specific ways to the human motor and auditory systems».

Hence, phonetics continued to be interested in describing speech sounds as a physical phenomenon (from acoustic, articulatory and auditory perspectives), whereas phonology developed increasingly formal apparatuses to describe the distinctive sounds used by language systems to build their words and the speaker’s mental representation of these sounds and words together with the rules controlling phonological processes (for a deep discussion of these issues cf. Durand & Laks 2002). Successive attempts advocated a sort of alliance of the traditional the approaches represented by phonetics and phonology (Lindblom 1986). However, Lindblom’s model did not represent an alliance that strive for a theory showing how phonological units receive different phonetic realizations (articulatory and acoustic) under different contexts, or how acoustical information can serve to reconstruct the phonological intentions responsible for their production (Bromberger & Halle 1986). The alliance proposed by Lindblom was essentially a physicalist alliance, which treats units of phonology as illusory by-products describable in articulatory and acoustic terms, so that phonological unit may be identifiable as phonetic ones.

Notwithstanding different scholars called for an *interface* (Blumstein 1991), an *interaction* (Keating 1991) or an *integration* (Ohala 1990) of the two disciplines (see also Pierrehumbert 1990), many phoneticians yet felt that the mental (abstract) entities posited in phonology are not subject to rigorous scientific investigation while phonologists argued that phonetics is a relatively uninteresting subfield of biology and physics, which is not useful to describe and explain certain linguistics aspects of human mind (with some notable exceptions as, for example, Archangeli & Pulleyblank 1994).

This mismatch has been crystallized into typical approaches which counterpose phonetics to phonology:

– Phonetics ended up being committed to describing/investigating pure acoustic, articulatory and perceptive properties of speech.
– Phonology is conversely involved in computational operations integrating the properties of speech into abstract (mental) representations subjected to categorical (discreet) processes.

### 1.2 The neurobiological perspective

While phoneticians and phonologists continued to separately investigate their subject matter, neuroscientists started exploring the phonetic and the phonological point of views from the neurobiological perspective. Indeed, the general issue of the neural representation of complex patterns is common to all neuroscience and has been investigated in many sensory modalities. In fact, if the spectrotemporal properties of speech are really converted into discrete (phonological) representations and these representations into appropriate motor commands to generate sequences of sounds, the brain is the unique responsible for the computational processes involved.

We have to note that the ability to categorize speech seems an ancient trait shared with many mammalian species. Studies on animals (chinchillas, gerbils, marmosets, ferrets, etc.) show clear evidence of categorical perception of speech sounds and conspecific vocalizations suggesting that the categorical processing is a general feature of auditory perception in vertebrates (Kuhl & Miller 1978; Ohl & Scheich 2005; Lu, Liang & Wang 2001; Mesgarani et al. 2008; Zoloth et al. 1979; Nelson & Marler 1989; see also Fitch 2010 for a discussion of these data). Also, animals vocalize modulating formant frequencies by changing the shape of their vocal tract: for instance, a cat may mew modulating formant frequencies to generate consonant-vowel-like ‘miaow’ sounds (Carterette et al. 1984).

Despite this shared property with animals, only humans have the ability to associate a finite sequence of sounds with potentially infinite concepts *signing* the external and internal world, and only humans can recursively combine sequences of signs (words) producing potentially infinite sentences. Moreover, the animal findings required massive training regimes in order to obtain categorical behavior patterns; conversely, human infants complete this task in the first stages of post-natal life (cf. Section 2.1). Crucially, animals do not need to convert these categories into representation structures that can make contact with higher-level levels of representations, e.g., morphemes, words, etc. Therefore, while the superficial behavior between the species may appear similar, the underlying computations required are likely to be quite different.

As suggested by Darwin (1871), the primary evolutionary changes required for human language were neural, not changes in vocal or auditory anatomy. What emerged as unique properties of the *Homo Sapiens* brain is the synchronization of neuronal activity along a functional cortico-thalamic network through a reentrant neural activity where neural information can be continuously interchanged between clusters of neurons: that is, for what concerns language computations, the synchronization of fronto-temporo-parietal clusters of neurons among themselves and with the thalamic nuclei (Edelman & Tononi 2000). Reentrant activity is present in all vertebrate brains but it is not synchronized with long-term and working memory. Reentrant activity per se is sufficient to generate primary conceptualization and categorization of the world (i.e., primary consciousness) but it cannot lead to effective learning processes. However, high-order conceptualization and categorizations are possible only when long-term memory may be synchronously integrated with working memory to result in continuous computational and representational processes (i.e., secondary consciousness) and then in learning (Edelman & Tononi 2000).

Accordingly, at one extremity of the speech perception process, in the inner ear, is the acoustic representation of speech, which, as we have seen, represents a general property of animal and humans. At the other extremity of the process are discrete (abstract) phonological representations, which can be manipulated by symbolic processes. Between these extremities there are multiple phonetic representations, which organize speech sounds into linguistically-relevant categories (Phillips 2001: 713-714). Thus, the analog representation of the acoustic of speech is converted in the digital representation of discrete (phonological) categories through the processing of phonetics categories, which are also analog, but which organize speech sounds into linguistically-relevant categories: the processing of the invariant properties of phonetic categories (i.e., distinctive features) leads to abstract representations.

In line with this perspective, the neural understanding of the high-resolution system for acoustic decoding and phonological encoding, tied to the ability for abstraction and an efficient memory mechanism, raises fundamental questions such as (i) whether spatially and temporally distributed activation patterns within the auditory cortex are directly implicated in speech sound processing; (ii) whether speech decoding is generated by pure bottom-up reflection of acoustic differences or whether they are additionally affected by top-down processes related to phonological categories and distinctive features. From this perspective, it seems that the phonetics-phonology relationship may be investigated according to Jacobson’s program unifying the two level of analysis within the neural patterns activations (cf. Manca & Grimaldi 2016).

### 1.3 Aims of the present work

After briefly introducing the different neurophysiological techniques used to investigate the auditory brain, I critically review the studies on speech perception (but also production data are taken into consideration). The findings of these studies are re-examined to understand how direct measurements of temporal and spatial brain activation can clarify the phonetics-phonology relationship. On the base of a recent work on speech processing computations and operations (Giraud & Poeppel 2012), I sketch a preliminary proposal aiming at explaining bottom-up and top-down processing. The idea is that discretization and phonological abstraction are the result of a continuous process that convert spectro-temporal states into neurophysiological states represented by nested cortical oscillatory rhythms spatially distributed in the auditory cortex.

## 2. Investigating the auditory brain

### 2.1 Techniques and methods

In auditory neuroscience, two techniques are widely used: electroencephalography (EEG) and magnetoencephalography (MEG), as they are the most powerful non-invasive tools with high temporal reliability (Roberts et al. 2000). Recently, also electrocorticography (ECoG) – an invasive approach used in clinical contexts where pre-surgical evaluation of cognitive processes is needed – is increasingly used to directly record auditory activity (Poeppel & Hickok 2015; Leonard & Chang 2016). MEG and EEG research into language processing is based on event-related potentials (ERPs) and event-related magnetic fields (ERMFs) recorded while the subjects are performing a cognitive task. Thereby stimuli are presented to subjects and markers are set into the EEG trace whenever a stimulus is presented. Then a short epoch of EEG/MEG around each marker is used to average all these segments. This is based on the logic that in each trial there is a systematic brain response to a stimulus. Practically, this means that one typically repeats a given experimental paradigm a number of times (say, >30 times), and then one averages the EEG/MEG recordings that are recorded time-locked to the experimental event.

However, this systematic response cannot be seen in the raw EEG, as there it is overlaid by unsystematic background activity (which is simply considered as noise). By averaging all the single epochs that are time-locked to the experimental event, only the systematic brain response should remain (i.e., those generate neural action potentials related to the stimuli), but the background EEG/MEG should approach zero (Sauseng & Klimesch 2008). The noise (which is assumed to be randomly distributed across trials) diminishes each time a trial is added to the average, while the signal (which is assumed to be stationary across trials), gradually emerges out of the noise as more trials are added to the average. These brain responses are named event-related potentials (ERPs) and event-related magnetic fields (ERMFs) reflecting the summated activity of network ensembles active during the task. ERPs/ERMFs are characterized by specific patterns called ‘waveforms’ (or ‘components’), which are elicited around 50-1000 ms starting from the onset of the stimulus and show positive (P) and negative (N) oscillatory amplitudes (i.e., voltage deflections). For instance, P100, N100, P200, P300, N400, P600 (or P1, N1, P2, and so on) are the principal components elicited during language processing starting from sound perception to semantic and syntactic operations. So, this technique provides millisecond-by-millisecond indices of brain functions and therefore provide excellent temporal resolution (Luck 2005).

The sensitivity to speech inputs in humans is very early. It seems that begins in the womb, when the auditory system of the fetus is matured and becomes attuned to a variety of features of the surrounding auditory environment (Partanen et al. 2013). In the first year of life, a clear perceptual transition from all the possible (universal) learning options to language-specific learning options emerges. Before 6-8 months of age, infants seem able to discriminate all the contrasts phonetically relevant in any of the world’s languages; by 12 months their discrimination sensitivity is warped by native phonemes while the perceptual sensitivity for non-native phonemes gradually declines (Werker & Tees 2005; Kuhl et al. 2006). According to Werker & Tees (1984), a recent neurophysiological study suggested that this cerebral reorganization around native categories is already formed at 6 months of age and may reflect a continuous process of neural commitment towards the first language categories (Ortiz-Mantilla et al. 2013).

### 2.2 The phonetic-phonological brain: acquisition and mapping auditory principles

The reshaping of the perceptual space in infants according to the phonetic properties of the mother tongue implies that constant computational processes on the signal are encoded online into abstract discrete representations of sounds by means of probabilistic and statistical operations computed by the brain on the acoustic signal (Kuhl 2004). A consequent hypothesis is that the acoustic-phonetic structures map directly onto clusters of neurons within the auditory cortex thanks to the specific sensitivity of nerve cells to the spectral properties of sounds: i.e., the so-called *tonotopic principle* (Romani et al. 1982). This coding of acoustic frequencies in different sites of auditory cortex is ensured by a selective activation process that begins early in the cochlear neurons regularly positioned along the basilar membrane (Moerel et al. 2014; Saenz, Langers 2014). Then, the neural signals emitted by cochlear neurons are transmitted in the brainstem and preserved up to the auditory cortex from the primary auditory cortex (A1) to the superior temporal gyrus (STG) and the superior temporal sulcus (STS) (Da Costa et al. 2011; Talavage et al. 2004). While pre-cortical processing seems to be common to all sounds, speech-specificity appears to arise at the cortex (Scott & Johnsrude 2003). Like retinotopy in vision, tonotopy is one of the most accepted models of cortical organization of the auditory pathway (Moerel et al. 2014) as also showed by studies on animals (Kaas & Hackett 2000; Rauschecker & Tian 2000; Mesgarani et al. 2008). Thus, from a linguistic perspective, speech-specific processing in the auditory cortex may be based on the *phonemotopy principle*.

In addition to the topographical separation of sounds of different frequencies, it has been suggested that latency of evoked responses may be a supplementary dimension for object encoding in the auditory system. Roberts & Poeppel (1996) demonstrated that there is a frequency dependence of latencies separate from stimulus intensity. Furthermore, recent animal data has shown that the precision of temporally based neural representations declines from periphery to the cortical regions entailing different encoding strategies for slow and fast acoustic modulations (Wang 2007). Thus, the temporal code may represent the ability of some pools of neurons to discharge at a particular phase of the structure of sounds (Zatorre & Belin 2001; Boemio et al. 2005). This temporal mechanism of auditory encoding is known as the *tonochrony* principle. That is, the latency of auditory evoked components appears to be sensitive to some stimulus properties; this suggests that the mechanism of tonochronic encoding might augment or supplement the tonotopic strategy in the frequency range critical to human speech (*phonemochrony*) (Roberts et al. 2000).

### 2.3 Event-related potentials, and event-related magnetic fields components

The auditory component widely investigated is N1, with its magnetic counterpart N1m, and mismatch negativity (MMN), with its magnetic counterpart MMNm. N1/N1m is a negative peak between 70 and 150 ms after the onset of an auditory stimulus (cf. Figure 2) that appears to be involved in the basic processing of speech sounds in auditory cortices (Woods 1995). It seems that the amplitudes and the latencies of the N1/N1m are relevant markers reflecting the cortical encoding of acoustic features of incoming speech sounds. The source location of the N1/N1m responses along the auditory planes seems to be driven by the spectral properties that are linguistically salient, e.g., the F1/F2 ratio for vowels, or the place of articulation for consonants, and then it may represent a good tool to investigate auditory cognitive processes.

**Fig. 2:**
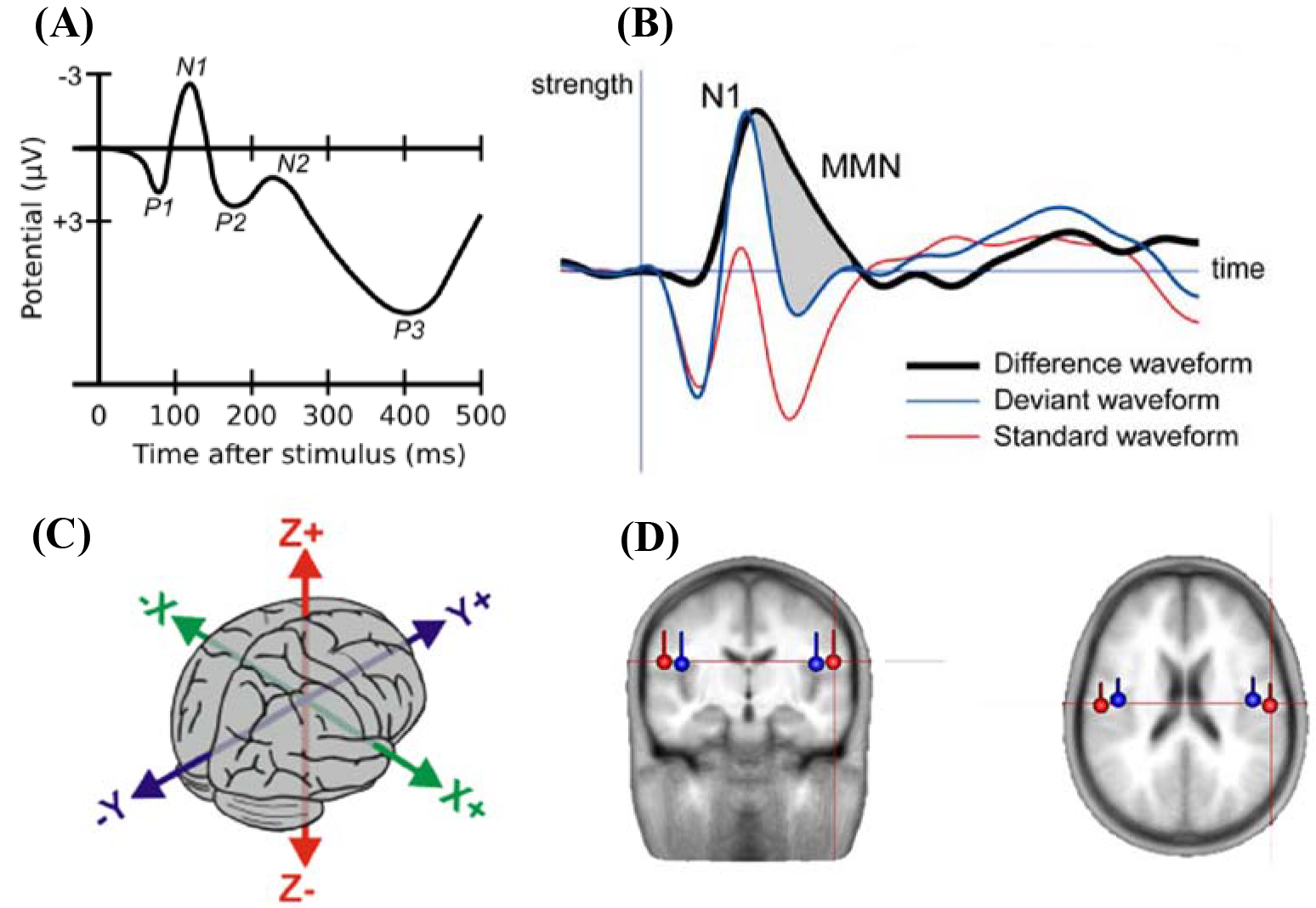
(A) Representation of the auditory N1 wave evoked from EEG to an auditory stimulus. The peak around 100 ms post-stimulus onset, measured in microvolts (μV) is evidenced (adapted from Lageman et al. 2012). (B) ERP waveforms evoked at a frontal scalp location by the standard and deviant sounds superimposed on the difference waveform in which the ERP to the standard has been subtracted from that to the deviant. The MMN appears as an enlarged negativity to the deviant sound as compared with the standard sound, following the N1 peak. Adapted from (Brattico 2006). (C) The 3D space within the brain along the classical Talairach coordinates: the direction of *x* axis is from left to right, that of *y* axis to the front, and the *z* axis thus points up. (D) Average location and orientation of the equivalent current dipole sources fitted in the bilateral auditory cortical areas. Adapted from Cirelli et al. (2014).

MMN/MNNm is a component temporally subsequent to the N1/N1m (cf. Figure 2), automatically and preattentively elicited by an acoustic change or by a rule violation between 150 and 250 ms post-stimulus onset (Näätänen 2001). Contrary to the N1/N1m, it is generated in a passive oddball paradigm, where subjects listen to frequent (standard) stimuli interspersed with infrequent (deviant) stimuli and attend to a secondary task (e.g., watching a silent movie). MMN/MMNm is visible by subtracting standard responses from deviant responses to the same acoustic stimuli and its amplitude seems to be directly correlated with the discriminability of the two stimuli involving both acoustic change-detection processes and phoneme-specific processes (Sussman et al. 2013). This component has been exploited to investigate (i) the categorical representation of phonemes in the subjects’ mother tongue (e.g. Näätänen et al. 1997); (ii) if the acoustic signal is mapped onto lexical representations through different levels of featural representation; in this case, N1m and MMNm have also been used together (Scharinger et al. 2011b; 2012), and (iii) if phonemic representations may eventually develop during second language acquisition (Grimaldi et al. 2014, and the literature therein).

As the signals measured on the scalp surface do not directly indicate the location of the active neurons in the brain, when interpreting EEG and MEG data, one has to solve the so-called the *inverse problem*: that is, the deduction of the source currents responsible for the externally measured fields measured on the scalp (Hallez et al. 2007). It is possible to simulate the neural activity by means of a dipolar model (Malmivuo et al. 1997). Dipoles are created by post-synaptic potentials of many single neurons oriented in the same direction and firing synchronously in response to the same event. Under stimulation, the dipoles from the individual neurons sum solving in a single equivalent current dipole (ECD) that seems to be the best approximation of auditory evoked potentials observed by sensors on the scalp. Location, orientation, and magnitude of the assumed ECDs provide information about the behavior of the activity under investigation (Luck 2005). The ECD can be described as a point located in a 3D space within the brain along the classical Talairach coordinates that represent the center of simultaneously active neural sources (Sanei & Chambers 2013): i.e., *x* (lateral-medial), *y* (anterior-posterior), and z (inferior-superior) axes (cf. Figure 2).

Do the available findings support a direct link between linguistic and neurophysiological primitives? That is, can tonotopy and tonochrony (as mirrored in N1m/N1 patterns) explain the properties of the phoneme computations and representations in terms of distinctive features within the auditory cortex?

#### 2.3.1 Amplitudes for vowels and consonants

From a general point of view, it has been shown that N1/N1m responses evoked by nonspeech tokens differ from those recorded with the speech tokens, which show stronger amplitude and longer latency. However, no indication of different underlying neural representations of speech sounds were found (Eulitz et al. 1995; Diesch et al. 1996; Poeppel et al. 1997; Swink & Stuart 2012). Subsequent works focusing on vowels discrimination tasks suggest that their representation is mainly guided by the spectral relations of frequencies rather than by abstract, phonological relevant features. As already showed for animals (Ohl & Scheich 1997), vowels with large F2-F1 distance (e.g., [i], [u]) elicited larger amplitudes than vowels with close F2-F1 formants peaks (e.g., [a]) (Diesch & Luce 1997; 2000; Obleser et al. 2003a; Shestakova et al. 2004) (cf. Fig. 1). These data have been interpreted at the light of the inhibition principle (Shamma 1985a: 1985b) according to which there exists a vowel-specific reduction of neuronal activity that depends on the vowel formant distance F2-F1 and that may be topographically organized along isofrequency contours.

All these previous studies used synthetic stimuli. When natural and large sets of vowels are compared, indications of phoneme distinction resulted broadly associated to the processing of featural variables. Scharinger et al. (2011a) and Grimaldi et al. (2016) investigated the entire Turkish and Salento vowel systems respectively. Turkish vowel system symmetrically distinguishes between high/non-high ([i, ⸀, y, u]/[⸀, ⸀, œ, ⸀]), unrounded front/back ([i, ⸀]/⸀, J]) and rounded front/back ([y, œ]/[u, ⸀]) vowels, while the Salento system is characterize by five vowels, i.e., [i, ε, a, ⸀, u], where [i, u] are high and [ε, a, ⸀] are non-high vowels. Both studies found that high vowels elicited larger amplitude than non-high vowels showing a categorical effect for phonological patterns. However, this result is also compatible with acoustic properties of speech sounds, as high vowels are significantly characterized by low F1 values, while low vowels by high F1 values (but see Monahan & Idsardi 2010, where a correlation of N1m with F1/F3 ratio is suggested). Note that Scharinger et al. (2011a) applied a mixed model statistical approach, testing whether the N1m complex was better accounted for by acoustic gradient predictors (acoustic model) or by distinctive features oppositions (feature model): interestingly, the feature model fitted the data better than the acoustic model suggesting that the processing of a vowel system may be relied on the abstract representation of articulatory plans and acoustic outputs, i.e., the binary opposition of distinctive features.

As for consonants, the data are scarce. Stable evidence pertains only to stops segments: stops ([b, p, d, t, g, k]) produce higher amplitudes than non-stop counterparts ([f], [r], [m], [r], [s]) (Gage et al. 1998). Also, the N1m amplitudes seem to vary as a function of the onset of the speech sounds with a higher amplitude for labial [ba] than alveolar [da] as compared to velar [ga] in both hemispheres. The overall difference in amplitude to stops vs. non-stops may be attributed to the acoustic differences in the onset dynamics of these two classes of stimuli. At the same time, it seems that N1m amplitude is sensitive to place of articulation. Within the class of stop consonants, the N1m peak amplitude resulted also sensitive to the feature Voicing. As revealed by Obleser et al. (2006), only intelligible voiced consonants ([d], [g]) yielded the stronger N1m amplitudes than their unvoiced counterparts ([t], [k]) did.

#### 2.3.2 Latencies for vowels and consonants

The N1m latency appears to be mainly related to the F1: i.e., high F1 values (e.g., [a] and [æ]) evoke shorter latency than low F1 values (e.g., [i] and [u]) (Diesch et al. 1996; Poeppel et al. 1997; Eulitz et al. 1995; Obleser et al. 2003a). Yet, works focusing on entire phonological systems highlight that the N1m/N1 changes seem related to the abstract processing of phonological features, although still tentatively: back vowels (e.g., [a, o, ⸀ u]) peaked later than the front vowels (e.g., [e, ε,ø i]) (Obleser et al. 2004a; 2004b; Grimaldi et al. 2016).

The mapping rules seem to proceed in a different way when testing the large set of the Turkish vowel system (Scharinger et al. 2011a). This study found that back vowels (e.g., [u]) were earlier than front vowels (e.g., [y]), and that the features Height and Round affected the timing neuronal strategies resulting in later responses to high (e.g., [i]) than non-high (e.g., [⸀]]) vowels and in faster N1m to unrounded vowels (e.g. [⸀]) than to the rounded counterparts (e.g., [u]).

The N1m latency was also found to be involved for point of articulation, Voice and the Back in naturally spoken syllables: the velar-back rounded CV syllable [go] elicited a later response than labial ([bø]), alveolar ([dø]), velar ([gø]) with front rounded vowel (Obleser et al. 2003b) confirming the critical role of point of articulation for temporal coding in human speech recognition (Roberts et al. 2000; Gage et al. 2002; Obleser et al. 2004b). According to the authors, this suggests that the assimilatory effect of a back vowel is very influential on a back consonant like [g]. Also, it has been observed that the N1m to isolated alveolar consonants [d] and [t] peaked earlier than responses to velar consonants [g] and [k], and voiced consonants [d] and [g] peaked later as compared to voiceless consonants [t] and [k] (Obleser et al. 2006). The authors, thus, proposed that the latency changes are mainly driven by the spectral (Place, spectral peak of [d-t] versus [g-k]) and temporal (Voicing, voice onset time, [d-g] versus [t-k]) features of the stimuli.

#### 2.3.3 Source generators for vowels and consonants

The dipole approach for modeling the N1m patterns and by observing the spatial arrangement into the brain assumes that the sound salient features for the phonological encoding drive the displacement of the N1m generators, which define specific arrangements (maps) on the cortical sheet. There are two perspectives of analysis concerning the spatial arrangement of speech sounds into the brain: the first observes the absolute distance of dipole generators in the cortical space (the auditory cortex in our case), the second calculates these distances relatively to a 3D space considering the anterior-posterior, lateral-medial, and inferior-superior Talairach axes (cf. Section 2, Fig. 1).

In line with the amplitude and latency findings, the absolute Euclidean distances between the representational centers of vowels reveal that the most dissimilar vowels in the F2-F1 space ([a-i]) tend to generate larger cortical distances than the most similar ones ([e-i]) (Obleser et al. 2003a; Mäkelä et al. 2003; Shestakova et al. 2004). However, some studies reported that vowels differing in only one place feature (front vowels [e]-[i] and front-rounded vowels [ø]-[y]) were closer than vowels maximally different for two or more features ([e]-[i] vs. back-rounded [o]-[u]) (Obleser et al. 2004b; see also Scharinger et al. 2011a).

The abstract representation of vowels emerges also for the relative distances along the Talairach axes. The N1m dipoles appear dependent on both spectro-temporal cues and phonetic features. The lateral-medial axis showed medial locations for vowels with F1 high frequencies (that is, low vowels) (Diesch & Luce 1997). Eulitz et al. (2004) found that the German vowel [i] with large spectral F2-F1 distance, was more medial than vowels with close formants peaks (e.g., [a]). Further studies described a broad cortical vowel distinction according to different phonological patterns. Obleser et al. (2004a) observed that the German back vowel [o] was lateral to the front vowel [ø]. On their part, Scharinger et al. (2011a) demonstrated that the dipole movements were more affected by phonological features than by acoustic properties and found a gradient for Round along this plane: i.e., rounded vowels ([y, œ, u, ⸀]) were at more lateral positions than unrounded vowels ([i, ⸀, ⸀, ⸀]).

The anterior-posterior plane seems responsive to the F1 and F2 values associated with Height and PoA. High vowels (e.g., [⸀] or [u]), with low F1, elicited a dipole that was located more anterior to the dipole of non-high vowels (a, [ɛ], [œ]), with high F1 values (Mäkelä et al. 2003; Scharinger et al. 2012). Also, the anterior-posterior plane seems correlated with the F2 values directly dependent by the Place phonological feature: back vowels (e.g., [⸀, ⸀, u, ⸀]) appeared at more posterior locations than front vowels (e.g., [i, e ⸀, y, œ]) (Obleser et al. 2004b; Scharinger et al. 2011a). As for consonants, fricative response was more posterior, on average, than the plosive response sources, the vowel and the nasal response sources (Kuriki et al. 1995). Obleser et al. (2003b) found that the differences in N1m source location were dependent on the point of articulation of the vowel but independent of the different syllable onsets: the front vowel [ø] elicited activity anterior to dorsal vowel [o]. Furthermore, the intelligibility alveolar [d] and [t] were more anterior than velar [k] and [g] irrespective of the voicing feature of the stimuli (Obleser et al. 2006). When labial and coronal consonants were compared (as in the couple [awa]-[aja] and [ava]-[a⸀a] respectively, labials elicited dipoles with more anterior locations than coronals (Scharinger et al. 2011b). This spatial location was independent of manner of articulation: anterior-posterior locations did not differ between [w] and [v] or [j] and [⸀]. A statistical model comparison showed that although the F2-F1 ratio was the best predictor for an acoustic model, a model built on the additional fixed effect place (labial/coronal) provided a better fit to the location data along the anterior/posterior axis. This might be interpreted as evidence for top-down categorical effects on the acoustically driven dipole location in auditory cortex.

Few studies reported significant results along the inferior-superior axis. Generally, low vowels resulted in superior location than high vowels (Shestakova et al. 2004; Eulitz et al. 2004; Obleser et al. 2003a). Conversely, Scharinger et al. (2012) revealed that the dipoles for the high [⸀] were approximately 7mm more superior to the dipoles for the low [æ], whereas the locations between [⸀] and [ε] and between [ε] and [æ] did not differ. Finally, Scharinger and colleagues (2011a) revealed a Round effect on the dipole locations, so that rounded vowels, which are acoustically marked by low F2 frequencies, were located at more inferior locations than dipoles to non-round vowels. However, when this effect was investigated for Front and Back vowels separately, the authors stated that the F1 and the related Height effects were, once again, the guiding rules for the cortical segregation within Front vowels only. For what concerns consonants, Obleser et al. (2006) showed front consonants (e.g., [d, t]) a more superior location than back counterparts ([k, g]).

#### 2.3.4 The MMNm/MMN

On the other side of the issue, also the MMNm/MMN studies offered important contributions on the understanding of categorical perception. Perception of vowels or VOT contrasts in the across-category conditions elicit MMNm/MMN amplitudes only for those segments having a contrastive role in the phonological system of listeners (e.g., Näätänen et al. 1997; Sharma & Dorman 2000). These results suggest that the MMNm/MMN is sensitive to the phonetic-phonological category distributions of the subjects’ native language (see Winkler et al. 1999; Grimaldi et al. 2014 for sensitivity to second language category). Also, studies on categorical discrimination (generally on consonant continua differing in the duration of VOT) highlighted that listeners are able to perceptually group the acoustic distinct tokens together to form a category. When they perceive a token from the other side of the category boundary, a change is detected as indexed by MMN (e.g., Sharma & Dorman, 1999; Phillips et al. 2000).

Phonemes used to contrastively distinguish lexical meaning may generate non-contrastive variants (i.e., allophones) that regularly appear in specific contexts because of the influence of adjacent vowels or consonants. Kazanina et al. (2006) used a multiple-token design with acoustic varying tokens for each of the stimuli to analyze the sound pair [t-d], which has allophonic status in Korean ([d] occurs between voiced sounds and [t] elsewhere) and a phonemic status in Russian. The results revealed an MMNm response for the Russian listeners but no response for the Korean listeners. The authors concluded that the phonemic representations, but not the allophonic ones, are computed from speech. These data, however, refer to VOT distinctions: what happens when vowels allophonic pairs generated by phonological processes are investigated? Miglietta, Grimaldi & Calabrese (2013) found different MMN patterns. They studied the allophonic variant generated by a phonological process (i.e., metaphony) characterizing southern Salento varieties that raises the stressed low-mid front vowel [⸀] to its high-mid counterpart [e] when followed by the unstressed high vowel [i]. MMNs were elicited for both the allophonic and phonemic conditions, but a shorter latency was observed for the phonemic vowel pair suggesting a rapid access to contrastive sound properties for the phonological patterns. Yet, the discrimination of the allophonic contrast indicates that also allophones – generated by specific rules of the grammar – are part of the knowledge of speakers and then of their memory representations. Thus, according to Calabrese (2012), the auditory cortex may have two ‘modes’ of perceiving speech: phonological perception (faster as it picks out from deeper language knowledge) and phonetic perception (slower as it refers to low-level acoustic properties only).

Finally, studies investigating whether phonemic representations in the lexicon are underspecified for non-contrastive distinctive features values in the language systems (Section 4.4) showed that MMNm/MMN are elicited only when the standard stimulus is fully specified for a distinctive feature while the deviant stimulus not (e.g., Eulitz et al., 2004; Scharinger et al., 2012). This is because a fully specified vowel (for instance for Height) in standard position should generate a strong expectation regarding tongue height specification that might be violated if the deviant to this standard sequence is an underspecified segment (for instance, a mid-vowel). This finding suggests that the MMNm/MMN may index more than just physical properties of the stimulus. However, Mitterer et al. (2003, 2006) found a symmetric MMN: a labial as deviant and a putatively underspecified alveolar as standard elicited the same MMN as the alveolar deviant with the specified labial standard. The same results were obtained by Bonte et al. (2005) with fricatives. Finally, evidence for underspecified features was not successively confirmed for labio-velar and palatal glides labial and palato-alveolar fricatives ([awa]-[aja] and [ava]-[a⸀a]) (Scharinger et al., 2011b).

### 2.4 The event related oscillatory rhythms perspective

Although the ERP approach has opened an important window on the time course and the neural basis of speech and language processing, more than 100 years after the initial discovery of EEG activity, researchers are turning back to reconsider another aspect of EEG, that is the event-related oscillations. This is because an increasing number of researchers began to realize that an ERP only represents a certain part of the event-related EEG signal. Actually, there is another aspect of extreme interest for the study of cognitive functions: the event-related fluctuations in rhythmic, oscillatory EEG/MEG activity. This view, indeed, might provide a new window on the dynamics of the coupling and uncoupling of functional networks involved in cognitive processing (Varela et al., 2001). In fact, substantial literature now indicates that some ERP features may arise from changes in the dynamics of ongoing EEG rhythms/oscillations of different frequency bands that reflect ongoing sensory and/or cognitive processes (Başar et al. 2001; Buzsáki 2006). More precisely, the EEG oscillations that are measured in a resting state become organized, amplified, and/or coupled during cognitive processes. It has been argued that ERP does not simply emerge from evoked, latency-fixed polarity responses that are additive to and independent of ongoing EEG (Sauseng et al., 2007): instead, evidence suggests that early ERP components are generated by a superposition of ongoing EEG oscillations that reset their phases in response to sensory input, (i.e., the external or internal stimuli generating cognitive activities). Therefore, event-related oscillations, further than to have the time-locked EEG information, permits the retrieval of non-phase locked EEG information related to the cognitive activity induced by the stimulus (cf. Figure 3(A) and (C)).

**Fig. 3:**
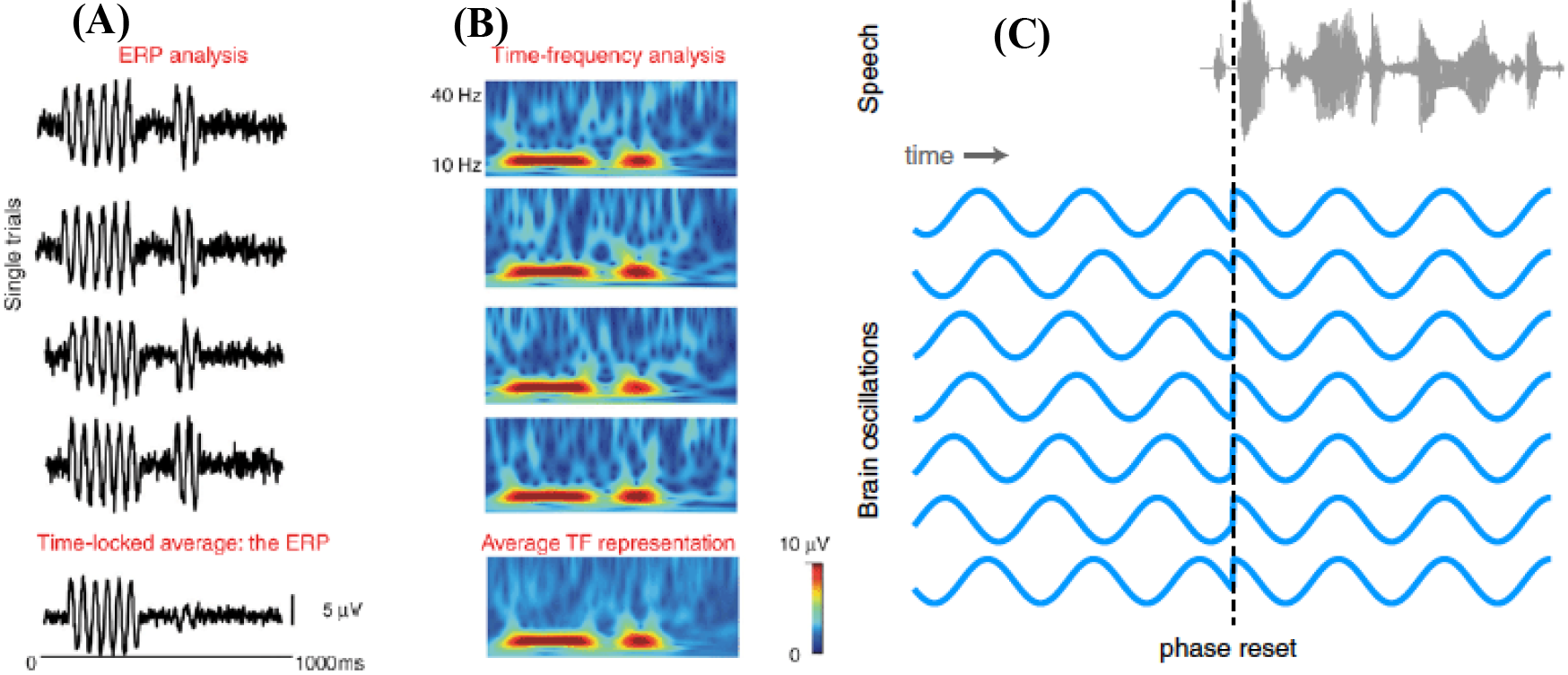
Simulated EEG data illustrating the difference between phase-locked (evoked) activity and nonphase-locked (induced) activity. (A): Single-trial EEG time courses showing two consecutive event-related responses (an amplitude increase at 10 Hz). The first response is phase-locked with respect to the reference time-point (t=0), and as a result this evoked response is adequately represented in the average ERP. The second response is time-locked, but not phase-locked to t=0, and as a result this induced response is largely lost in the average ERP. (B): time-frequency (TF) representations of each single trial, with red colors coding for the amplitude increase at 10 Hz. Crucially, the average TF representation contains both the phase-locked and the non-phase-locked responses. Adapted from Bastiaansen, Mazaheri & Jensen (2012). (C) Illustration of the phase of ongoing neural oscillations being reset by an external stimulus. Prior to the stimulus event the phases of the oscillations are random, but following the stimulus they are aligned. Adapted from Pelle & Davis (2012).

Within this perspective, ongoing cerebral activity can no longer be thought of as just relatively random background noise (the non-phase EEG activity), but as a whole containing crucial information on the dynamical activity of neural networks: thus, the EEG and ERP are the same neuronal event, as the ERP is generated because of stimulus-evoked phase perturbations in the ongoing EEG. A fundamental feature of the phase-resetting hypothesis is that following the presentation of a stimulus, the phases of ongoing EEG rhythms are shifted to lock to the stimulus. From this, it follows that during pre-stimulus intervals, the distribution of the phase at each EEG frequency would be random, whereas upon stimulus presentation, the phases would be set (or reset) to specific values (for each frequency). The resetting of the phases causes an ERP waveform to appear in the average in the form of an event-related oscillation (Makeig et al. 2002; Penny et al. 2002; Klimesch et al. 2004).

Unlike ERP (based on the analysis of components), event-related oscillations are based on the time-frequency analyses (e.g., Gross, 2014). One such method is wavelet analysis. The general idea is that not all relevant EEG activity is strictly phase-locked (or evoked) to the event of interest (Buszáki, 2006). Obviously, this activity shortly before stimulus onset is mostly not visible in ERPs due to cancellation; nevertheless, this pre-stimulus baseline activity may have a crucial impact on the observed ERPs (Klimesch, 2011). Time-frequency analyses enable us to determine the presence of oscillatory patterns in different frequency bands over time. Thus, with wavelet analyses, it can be established whether oscillatory activity in a specific frequency band, often expressed in power (squared amplitude), increases or decreases relative to a certain event, as represented in Figure 3(B).

The importance in considering the non-phase locked event-related oscillations consists in the fact that, contrary to phase-locked responses as ERPs, they reflect the extent to which the underlying neuronal activity synchronizes. Synchronization and de-synchronization are related to the coupling and uncoupling of functional networks in cortical and subcortical areas of the brain (Varela et al., 2001; Bastiaansen, Mazaheri & Jensen 2012). This aspect, of course, is related to how different types of information, which are stored in different parts of the network, are integrated during computational and representational processes. Importantly, elements pertaining to one and the same functional network are identifiable as such by the fact that they fire synchronously at a given frequency (cf. Figure 3). This frequency specificity allows the same neuron (or neuronal pool) to participate at different times in different representations. Hence, synchronous oscillations in a wide range of frequencies are considered to play a crucial role in linking areas that are part of the same functional network. Importantly, in addition to recruiting all the relevant network elements, oscillatory neuronal synchrony serves to bind together the information represented in the different elements (Gray et al. 1989).

In brief, the ERP approach is based on time domain analysis and wants to know: when do things (amplitudes and latencies) happen? The oscillatory approach is based on Frequency domain (spectral) analysis (Fourier analysis), analyzes magnitudes and frequencies of wave (renouncing to time information) and wants to know: when do which frequencies occur and when their power increases/decreases? That is, the ERP perspective treats peaks and troughs as single events, while the oscillatory approach as separate entities.

Also, recent findings suggest that ERP components and oscillatory rhythms represent complementary measures of the neural processing underlying high-level cognitive processing: the extent at which they capture same aspects of language processing is at the moment unclear (Wang 2010). Actually, neural oscillations in the low delta-theta ranges and high beta-gamma ranges have been shown to temporally correlate with different speech and categorial processes (including lexical processing) (Kösem, van Wassenhove 2017).

#### 2.4.1 The event related oscillatory rhythms data

The idea being investigating oscillatory rhythms is that endogenous fluctuations in neural excitability can be controlled by the brain so that they may be entrained (synchronized) with external rhythmic (and predictable) inputs as speech sounds (Zoefel, VanRullen 2015). In fact, the frequency of the speech envelope (roughly defined as the sum of energy across sound frequencies at a given point in time) is relatively stable between 2 and 8 Hz and phases of low phonetic information rhythmically alternate with phases of high phonetic information. It seems that there is a remarkable correspondence between average durations of speech units and the frequency ranges of cortical oscillations: phonetic properties (duration of 20-50 ms) are associated with gamma (>50 Hz) and beta (15-30 Hz) oscillations; syllables, and words (mean duration of 250 ms) with theta (4-8 Hz) oscillations; sequences of syllables and words embedded within a prosodic phrase (500-2000 ms) with delta oscillations (<3 Hz) (Ghitza 2011). In brief, there would exist a principled relation between the time scales present in speech and the time constants underlying neuronal cortical oscillations that is both a reflection of and the means by which the brain converts speech rhythms into phonemes (Giraud, Poeppel 2012). The same question posited for ERPs components arises: is the oscillatory rhythms entrainment a generic processing of acoustic patterns or is it informative of specific categorial processing in speech? (see Kösem, van Wassenhove 2017 for a critical review).

High-frequency gamma rhythms are involved in speech perception. A recent ECoG study succeeded at reconstructing the original speech input (series of words and pseudowords) by using a combination of linear and nonlinear methods to decode neural responses from high-gamma (70-150 Hz) activity recorded in auditory cortical regions. The decoded speech representations allowed readout and identification of individual words directly from brain activity during single trial sound presentations (Pasley et al. 2012). Mesgarani et al. (2014) analyzed the same high gamma cortical surface field potentials, which correlate with neuronal spiking, while participants listened to English natural speech samples (500 sentences spoken by 400 people) and the neural answers were recorded with ECoG. Most speech-responsive sites were found in posterior and middle STG. Neural responses showed distributed spatiotemporal patterns evoked during listening in line with different subsets phonetic-phonological patterns. Thus, the ECoG electrodes collocated in the auditory cortex divide STG sites into two distinct spatial gradients: obstruent- and sonorant-selective neural sites. The obstruent-selective group resulted divided into two subgroups: plosive and fricative electrodes. Among plosive electrodes, some were responsive to all plosives, whereas others were selective to place of articulation (dorsal /g/ and /k/ versus coronal /d/ and /t/ versus labial /p/ and /b/) and voicing (separating voiced /b/, /d/, and /g/ from unvoiced /p/, /t/, and /k/). Fricative-selective electrodes showed weak, overlapping selectivity to coronal plosives (/d/ and /t/). Sonorant selective cortical sites, in contrast, were partitioned into four partially overlapping groups: low-back vowels, low-front vowels, high-front vowels, and nasals. Although very interesting, this neural mapping showed just a selective response to subsets of phonemes. Also, the authors found selectivity for some higher-order acoustic parameters, such as examples of nonlinear, spatial encoding of VOT, which could have important implications for the categorical representation of this temporal cue. Furthermore, they observed a joint differential encoding of F1 and F2 at single cortical sites, suggesting evidence of spectral integration processing.

Interestingly, high-gamma oscillations play a crucial role also in speech production. Bouchard et al. (2013) found that these rhythms are generated in the ventral sensory motor cortex – a cortical region that controls the vocal articulators – during the production of American English vowels and consonants. Populations of neurons in this cortex showed convergent and divergent dynamics during the production of different phonetic features. The dynamics of individual phonemes were superimposed on a slower oscillation that characterizes the transition between consonants and vowels. Although trajectories were found to originate or terminate in different regions, they consistently pass through the same (target) region of the state-space for shared phonetic features. Consonants and vowels occupy distinct regions of the cortical state-space. Large state-space distances between consonant and vowel representations may explain why it is more common in speech errors to substitute consonants with one another, and vowels with vowels, but very rarely consonants with vowels or vowels with consonants (that is, in ‘slips of the tongue’). A recent study, Rapela (2016), confirmed this view by showing that rhythmic speech production (i.e., sequence of syllables) modulates the power of high-gamma oscillations over the ventral sensory motor cortex and the power of beta oscillations over the auditory cortex (due to the auditory feedback necessary control acoustic-articulatory outputs). He found significant coupling between the phase of brain oscillations at the frequency of speech production and their amplitude in the high-gamma range (i.e., phase-amplitude coupling). Furthermore, the data showed that brain oscillations at the frequency of speech production were organized as traveling waves and synchronized to the rhythm of speech production.

Additional evidence suggests that high frequency activity tags the timing of abstract linguistic speech units and structures (Kösem & van Wassenhove 2017). For instance, high-frequency activity has been shown to reflect the result of top-down word parsing: high-frequency oscillatory power dynamics systematically delineated the boundaries of perceived monosyllabic words in ambiguous speech streams (Kösem et al. 2016). High-frequency activity also represented, concurrently with delta oscillations, the segmentation of longer abstract linguistic structures (phrases and sentences) in continuous speech (Ding et al., 2016).

Di Liberto, O’Sullivan, Lalor (2015) performed an interesting EEG experiment in order to provide evidence of the neural indexing of the categorical perception of phonemes using series of natural sentences. They adopted a cross-validation approach to quantify how well each speech representation related to the neural data testing five speech representations: (1) broadband amplitude envelope; (2) spectrogram; (3) time-aligned sequence of phonemes; (4) time-aligned sequence of phonetic features (according to Chomsky & Halle 1968); and (5) a combination of time-aligned phonetic features and spectrogram. Crucially, the combined time-aligned phonetic features and spectrogram model of representation outperformed all other models principally in the theta but also in the delta bands. Within latencies of 150-200 ms vowels and consonants resulted neurally mapped on hierarchical clusters according to different acoustic-articulatory features. As the experimental subjects were exposed to sequences of natural sentences (derived from a professional audio-book version of a popular mid-20th century American work of fiction), one can suppose that theta oscillations are involved in phonological discretization processes, while delta in suprasegmental processing.

Accordingly, theta (3-8 Hz) oscillations are also recruited during phonemic restoration (Riecke et al. 2012; Sunami et al. 2013; Strauß et al. 2014). It has been showed that theta oscillations may be involved in the identifications of consonants in syllables ([da], [ga]) matching the different temporal onset delays in natural audiovisual speech between mouth movements and speech sounds (Ten Oever & Sack 2015; see also Kösem et al. 2016).

From a general point of view, delta oscillations have been associated with the processing of non-speech-specific attentional and predictive modulations of auditory processing (Kösem et al. 2016), but they do not serve the same purpose as theta oscillations. Recent findings suggest that they can play a crucial role in parsing speech units in the acoustic signal (Buiatti et al. 2009). According to Ghitza (2011) this rhythm plays an important role in prosodic parsing, which pertains to sequences of syllables and words, hence tapping contextual effects: as such, the delta oscillator may interact with the theta, beta, and gamma oscillators in a top-down fashion. Recently, Ding et al. (2016) the observed delta frequency corresponded to the syntactic complexity of the heard speech signal. Series of phrases constituted of two monosyllabic words were associated with a neural entrainment at 2 Hz, and sentences composed of four monosyllabic words induced a delta response at 1 Hz. These data suggest that delta oscillations could also be involved in the combination (or unification) of word units into a longer and more abstract linguistic structure, that is in the full range of hierarchical linguistic structures from syllables to phrases and sentences. Along these lines, Murphy (2016) proposes a revolutionary theory of neurolinguistics, proposing that nested oscillations execute elementary linguistic computations involved in phrase structure building. Murphy’s theory goes considerably beyond existing models, and it will be of interest for future research whether causal-explanatory power can also be attributed to nested oscillations with respect to the phonological system.

In brief, nested low theta and high gamma frequency activity may be associated with the output of phonetic-phonological processing, while delta phase resetting and entrainment may reflect the combinatorial processes underlying sentence unification, including phrase level prosody processing.

It is important to note that it is impossible to assign a single function to a given type of oscillatory activity (Başar et al. 2001). It is thus unlikely that, for instance, theta has a single role in language processing. In fact, theta’s role and its varying patterns of coherence as a function of task demands may be better seen in its relationship to beta and gamma (and same thing is true for the other oscillatory rhythms). It is arguable that the same oscillatory rhythms play different roles in speech and language processing (as in other cognitive processes) with simultaneous changes in the coherence patterns in the different frequency ranges. Actually, theta rhythms seem involved also in retrieving lexical semantic information and controlling processes with multiple items; on the other hand, gamma-band neuronal synchronization is involved in sentence-level semantic unification; alpha phase acts not only in decisional weighting, but also in semantic orientation, in creative thinking, and lexical decisions; beta synchronization serves to bind distributed sets of neurons into a coherent representation of (memorized) contents during language processing, and, in particular, building syntactic structures (cf. Grimaldi in press for a detailed discussion).

## 3. What does neurophysiological data tell us?

Cumulatively, the reviewed N1m/N1 studies suggest that the acoustic-articulatory patterns of speech sounds affect amplitudes, latencies and the spatial arrangement of neural sources. However, finding a ubiquitous system for speech-specific processing in line with the phonological features hypothesis is difficult. When we look at the data, it is hard to disambiguate between N1m/N1 evidence suggesting pure acoustic patterns and those indicating abstract phonological features. Solid evidence is that the acoustic distance between the first two formants of vowels is preserved in the auditory cortex and is directly detectable in sensor and source data (as found in the mammalian auditory cortex). This implies neuronal sensitivity interactions between spectral components during vowel discrimination that does not necessarily require separate formant representations (in the sense of feature extraction).

At the same time, these data coexist with further findings suggesting that acoustic-articulatory properties are affected by top-down features processing such as Height, Place and Round (this seems true also for Voice and Place for consonants), as showed by amplitudes, latencies and Talairach spatial coordinates in the auditory cortex. Note, however, that also when an entire vowel space of a language (Turkish) has been investigated (Scharinger et al. 2011) and evidence for the spatial arrangements of neural sources according to distinctive features seem strong, it is very difficult to disentangle the effect of acoustic parameters (F1, F2, F3) from that of distinctive features in neural maps processing and representation. In fact, the comparison of the source locations suggests that the auditory spatial encoding scheme is based both on acoustic bottom-up and top-down phonological information, and only a sophisticated statistical analysis of the data led to suggest that the continuous acoustic parameters per se do not warrant the understanding of abstract featural representation. But, again, it remains unclear how the correlation between acoustic parameters (e.g., F1) and feature patterns (tongue height) are eventually processed in different neural dimensions in perceiving sounds of speech.

To sum up, while the N1m/N1 findings show that the electrophysiological sensitivity to the properties of the stimuli is not exclusively correlated with their physical attributes, but rather with top-down processes associated with phonological categories, they are not so strong to preclude purely acoustic explanations of the auditory activity involved in speech processing: the amplitude data are not sufficient to prove a correlation with phonological patterns and the latency results appear contradictory (cf. Section 4). For example, while Obleser et al. (2004a, 2004b) and Grimaldi et al. (2016) showed that back vowels peaked later than non-back vowels, Scharinger et al. (2011a) revealed the reverse pattern. Finally, only one study with consonants has obtained strong evidence of latency modulations for Place and Voice (Obleser et al., 2006). Although it seems that latencies are probably correlated also with Height and Round features (Scharinger et al., 2011), it is difficult to establish to what extent the activity involved in speech encoding reflects merely mechanisms of spectro-temporal information extraction or rather of top-down processes related to phonological computations.

On the other side of the issue, also the MMNm/MMN studies seem to support this view. While many studies have provided neural evidence for categorical (and then abstract) representations of speech sounds and that distinctive features may be the neural signature of perception mechanisms, others suggest that also sensory inputs guide speech computation and representation. A crucial data for this perspective is represented by the fact that allophones generated by a phonological process (that is, a rule of the grammar) elicited the same MMN of phonemes (with the later characterized by an early neural answer): this suggests that both acoustic and abstract modes are immanent in the mapping of auditory inputs into higher perceptual representations and that they are related to memory processes (i.e., the listeners’ knowledge of the phonological system).

In principle, this difficulty in showing the abstract representation of speech sounds in terms of neural processing may be due to different factors, as the nature of the stimuli, the experimental procedures developed, etc. However, I suggest that the constant joined effect of acoustic parameters and phonological processing in neural maps representation may reveal important properties of the phonetics-phonology relationship rather than showing the limits of these studies.

The oscillatory approach permits us to address this issue from another perspective and to clarify the phonetics-phonology relationship from a new viewpoint. The available data (not yet extensive) suggest a neural multidimensional feature space for speech perception and production encoding and not precise unit boundaries for each speech sound feature (cf. Mesgarani et al. 2014 and Bouchard et al. 2013). This flexible encoding of neural patterns may account for coarticulation and temporal overlap over phonemes. Thus, responses to group of acoustic parameters show a neural selectivity for specific phonetic features dynamically encoded by oscillatory rhythms: this selectivity emerges from combined neural tuning to multiple acoustic parameters that delimit phonetic contrasts. If we assume that the auditory system has evolutionally become tuned to the complex acoustic signal rhythmically produced by vocal apparatus, it is possible to hypothesize that neural oscillation patterns reflect the patterns of decoded speech (Giraud, Poeppel 2012). In other words, the input signal generates phase resetting of the intrinsic oscillation in auditory cortex in line with the spectro-temporal properties of speech stimuli (Schroeder & Lakatos 2009). As gamma oscillations have a period of approximately 25 ms, they may provide 10-15-ms window for integrating spectrotemporal information (low spiking rate) followed by a 10-15-ms window for propagating the output (high spiking rate). However, because the average length of a phoneme is about 50 ms, a 10-15-ms window might be too short for integrating this information (Arnal, Poeppel & Giraud 2016). Accordingly, thanks to computational models of gamma oscillations, it has been suggested that a 50-ms phoneme could be correctly processed with two/three gamma cycles (Borgers et al. 2005; Shamir et al. 2009).

So, the temporally limited capacity of gamma oscillations to integrate information over time possibly imposes a lower limit to the phoneme length. On the other hand, the average length of syllables is approximately 150-300 ms: because the theta band (4-8 Hz) falls around the mean syllabic rate of speech (~ 5 Hz), the phase of theta-band activity likely tracks syllabic-level features of speech. In short, it is arguable that the auditory cortex uses gamma oscillations to integrate the speech auditory stream at the phonemic timescale, and theta oscillations to signal syllable boundaries and orchestrate gamma activity (Ghitza 2012; Arnal, Poeppel, Giraud 2016). The gamma and theta bands, which concurrently process stimulus information, lie in a nesting relation such that the phase of theta shapes the properties (amplitude, and possibly phase) of gamma bands.

## 4. A preliminary proposal to explain bottom-up and top-down processes

### 4.1 Analog and digital processing of speech sounds

How the data until now discussed can explain the classical linguistic assumption that, starting from the acoustic signal, the phonetics properties are analyzed online and decoded into abstract representations (distinctive features)? That is, how neural data can explain the notion that speech perception is discrete (categorical) (Repp 1984; Scott & Evans 2010; Liebenthal & Bernstein 2017; see also Van Rullen & Koch 2003)? It is well known that categorical perception relates to the way in which a continuous sequence of equal physical changes in a stimulus is perceived and represented as different categories of stimuli. According to Giraud & Poeppel (2012)’s model, an incoming speech signal generates the neuronal (spiking) activity in the form of oscillatory rhythms (note that spike, or action potential, is a neuron discharged signal in response to stimuli, generally called synapse). The activity in the gamma band has a tightly coupled relation to spike trains, regulating spike patterns. Finally, neuronal excitability is modulated such that acoustic structure of the input is aligned with neuronal excitability. By this hypothesis, the theta and gamma oscillations act (i) by discretizing (sampling) the input spike trains to generate elementary units of the appropriate temporal granularity for subsequent processing and (ii) by creating packages of spike trains and excitability cycles.

Giraud & Poeppel (2012) argue that a functional consequence of the modulation of low and high pyramidal neurons by theta and gamma oscillators is the organization of spike timing and the ensuing discretization of the cortical output. Spike discretization serves to present the stimulus in discrete chunks (segments) from which many different types of computations can be performed thanks to a principled relation between the time scales present in speech and the time constants underlying neuronal cortical oscillations. Ultimately, the process permits phonological abstraction, generating discrete representations that make contact with spatially distributed phonemic and syllable representations underlying recognition (Giraud & Poeppel 2012: 3). This view, as clearly suggested by Arnal, Giraud & Poeppel (2016: 466), presupposes that the continuous stream of natural speech is analyzed online and decoded by discontinuous, separable modes in the auditory cortex (i.e., neural spikes) in a sort of binary code: for each oscillatory cycle, output neurons may fire or not fire, which constitutes a binary code reflecting the shape of the speech envelope (Giraud & Poeppel 2012: 4).

However, as observed by Obleser, Herrmann & Henry (2012), the time scales present in speech cross functional boundaries between oscillatory bands in the human brain. Furthermore, while there are studies suggesting that synaptic activity onto pyramidal neurons (for instance in the hippocampus) is altered in discrete steps (Petersen et al. 1998; O’Connor et al. 2005), other studies show that a property of synapses is to change continuously, also for what concerns memory processes (Tanaka et al., 2008; see also Chaudhuri & Fiete 2016; Sjöström & Gerstner 2017). Although a longstanding tradition has hypothesized that information in a spike train is digitally (i.e., discreetly) encoded by the number of spikes in a given time interval, other empirical and theoretical perspectives suggest that information can also be encoded in the precise timings between single spikes as they arrive, namely through analog encoding (Rieke et al. 1999). Recently, Mochizuki & Shinomoto (2014) confirmed this perspective showing that fractions of neurons in subcortical regions (e.g., thalamic nuclei) exhibit digital patterns while cortical regions (e.g., primary visual cortex and middle temporal area) exhibit analog patterns. Therefore, it seems that both analog and digital forms are used by neurons to encode that is sent to and from the brain.

### 4.2 From spectro-temporal states to neurophysiological states through analog and digital spike-trains

I hypothesize that speech discretization is not due only to rhythmic packages of spike trains as resulted by the precise alignment of acoustic structure with neuronal excitability. Instead, discretization and phonological abstraction are the result of a continuous process which converts physical states into other physical states. I hypothesize that the *spectro-temporal states* are continuously converted into the *neurophysiological states* so that properties of the former states undergo changes interacting with the latter states until a new equilibrium is reached: such equilibrium is represented by synapses spatially distributed within the auditory cortex by means synchronized activities of oscillatory rhythms. At the production level, the inverse dynamical and continuous conversion happens: i.e., the neurophysiological states are converted into the spectrotemporal states through the same principle (here, however, I will restrict my hypothesis at the perception level only).

Why might the cortical brain use continuous process to convert the spectro-temporal states into the neurophysiological states rather than into well-separated discrete states? Two steps are needed to encode a continuous variable in a set of well-separated discrete stable states in some other coding dimension: (i) discretizing, and then (ii) choosing how to map such discrete values. In general, there is no metric-preserving mapping and the complexity of such encoding process is considerable. Conversely, the values of a continuous variable can be naturally and continuously mapped onto (quasi-)continuous states, preserving metric relationships between different values of the variable. The encoding is relatively simple, with the selection of a different storage state for a different value based on the change in the variable value (cf. Chaudhuri & Fiete 2016).

I suggest that a continuous process is better suitable for learning and memory processes. Mapping spectro-acoustic variable onto a continuous multiple of matching neural states can preserve much more information useful for computational and memory processes. With a neural representation that preserves the information of the external variable input, it becomes possible, given an input with spectro-temporal variable patterns, to directly update the neural states according to the values of the external variables. For instance, when a learner is exposed to a linguistic system he/she does not only have to acquire which sounds represent phonemic categories but also to acquire phonological processes which generate allophonic variants in the appropriate environments (cf. Miglietta, Grimaldi & Calabrese 2013). As we have seen (cf. Section 3), a clear perceptual transition from the universal learning options to language-specific learning options emerge very early in infants: this view is compatible with a model of phonology in which the acquisition of phonemic categories parallel occurs with the learning of phonetic distribution patterns and their relationships. Moreover, a subsequent process is required to map the abstract representation of speech sound onto concepts, so as to generate lexical items, labelling, morphological computations, and Spell-Out transfer operations to the conceptual-intentional and sensory-motor interfaces, and ultimately to have a coherent representation of sentences. This process must be necessarily continuous, probably controlled by theta-gamma, alpha-gamma-beta and gamma-beta-theta nested oscillations (see Grimaldi in press).

The process that convert spectro-temporal states into neurophysiological states starts from the cochlea response where different frequencies reflect spectro-temporal properties of vowels and consonants, and continue through the auditory pathways where these frequencies are progressively converted into oscillatory rhythm in the auditory cortex creating nested neural oscillatory states resulting in distinctive features representations. As in the cochlea a tonotopic coding of acoustic frequencies is ensured by the selective activation of the cochlear neurons regularly positioned along the basilar membrane (Moerel et al. 2014; Saenz & Langers 2014), also the auditory cortex ensures a spatiotemporal mechanism as a tool to discretize speech sounds (the so-called tonotopic principle: Romani et al. 1982). In production, auditory and motor cortices integrate their activity through the oscillatory rhythms in order to produce the acoustic output.

I hypothesize that, from the cochlea, speech sounds activate a preliminary spectrotemporal computation in the auditory thalamus, or medial geniculate body (cf. Bartlett 2013). Here, the first conversion of spectrotemporal states in neurophysiological states happens, and neurons encode the signal in digital form, so as to phase-lock to speech envelope modulating theta oscillations. In this state, phonetic categories are broadly mapped. After reset, theta oscillations generate a nested modulation of gamma oscillations in the primary auditory cortex where neurons begin to decode the signal into continuous (analog) linguistically-relevant categories. Finally, the last neurophysiological state takes place in secondary auditory areas (superior temporal gyrus, Brodmann area 22), where invariant properties of phonetic categories (i.e., distinctive features) are mapped leading to abstract representation through analog and spatially distributed gamma oscillations (cf. Figure 4).

**Fig. 4:**
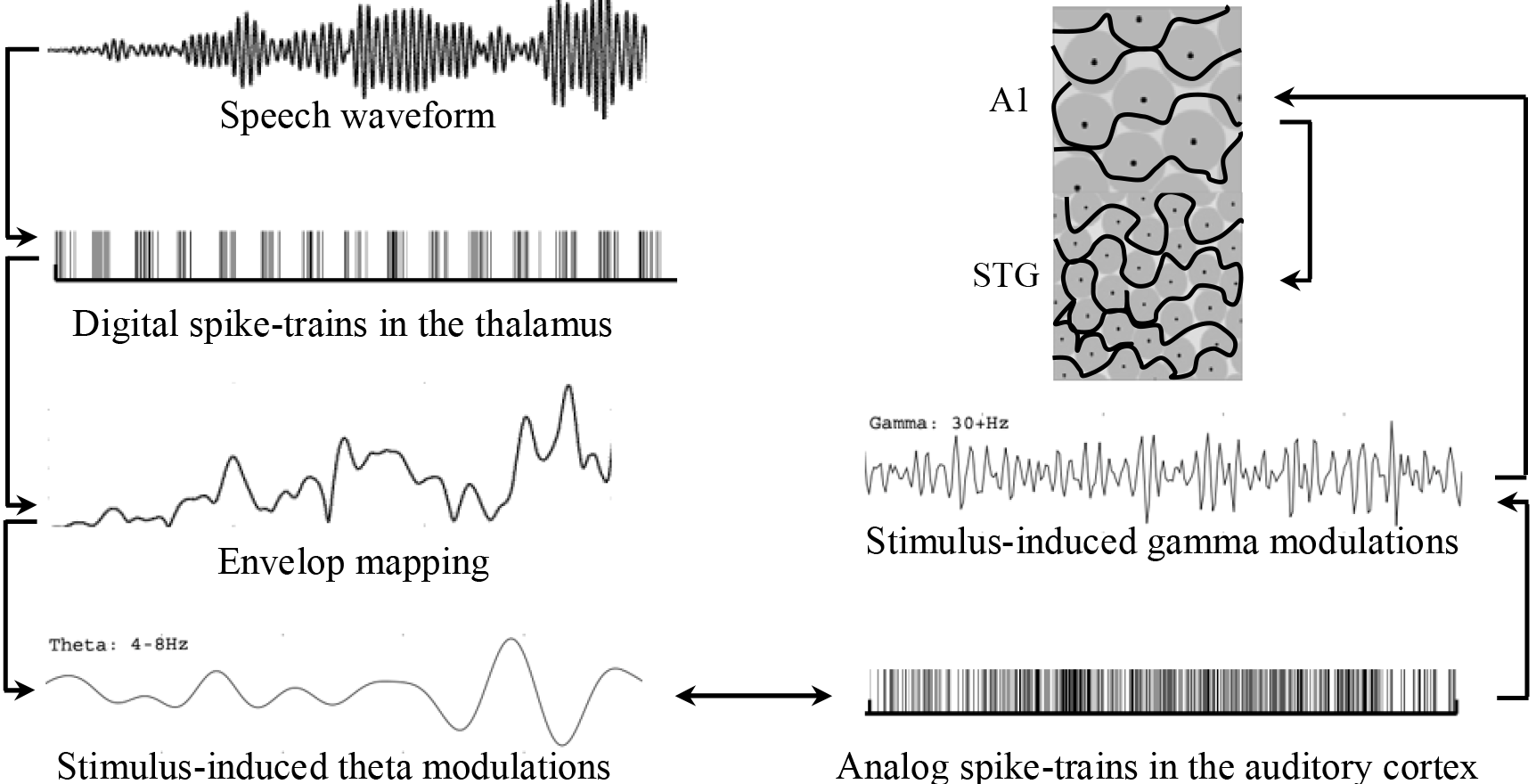
a model of speech perception based on digital and analog spike-trains processing that convert spectro-temporal states within neurophysiological states. Acoustic stimuli induced digital spike-trains in the thalamus (medial geniculate body) and theta oscillations are modulated to map speech envelop. Theta oscillations are coupled with gamma oscillations and analog spike-trains decode broad phonetic categories within the primary auditory cortex (A1) and invariant properties of phonetic categories (distinctive features) within the superior temporal gyrus (STG, Brodmann area 22).

**Fig. 6:**
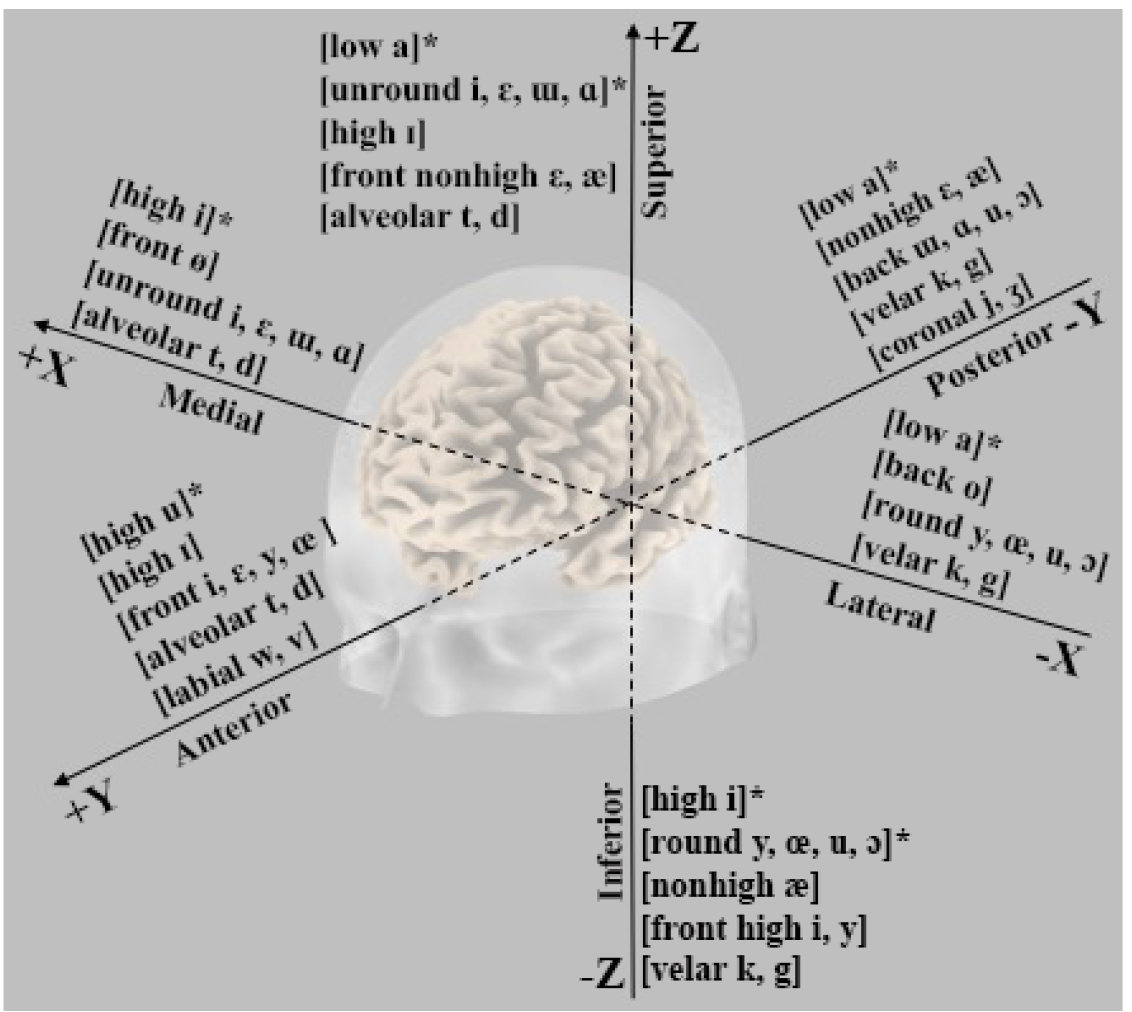
Graphical representation of the main trends emerging from the N1m ECD analysis along the Talairach three-dimensional spaces slicing human brain in lateral-medial (*x*), anterior-posterior (*y*), and inferior-superior axis (*z*). The symbol (*) indicates that the topographical gradient was explained in terms of acoustics effects rather than of featural variables.

I assume that, at the end of the process, phonological segments (and then categorical processes) take the form of continuous neural states represented by nested cortical oscillatory rhythms spatially distributed in the auditory cortex (as suggested by Bouchard et al. 2013; Mesgarani et al. 2014). Within this perspective, distinctive features (i.e., the relevant representational linguistic primitives) are represented by both spatially local and distributed neural selectivity (Mesgarani et al. 2014; Di Liberto et al. 2015). So, the abstract representation of distinctive feature is reached when nested theta and gamma bands (and probably also beta bands: cf. Ghitza 2011) reach the maximum power within the appropriate auditory space tonotopically deputed to transform the peculiar properties of a speech stimulus into discrete neural states (i.e., distinctive features). While Giraud & Poeppel (2012) see the spatially distributed phonemic and syllable representations as a consequence of discrete chunks generated by spike trains, I suggest that discretization processes emerge at the cortical level when different neurophysiological states result spatially distributed within the secondary auditory cortex, without the need for the alignment of acoustic structure of the input with neuronal excitability. Therefore, the binary properties of distinctive features that permit polar oppositions is neurally instantiated in the tonotopic presence or absence of appropriate neurophysiological states within the secondary auditory cortex. In brief, discretization and categorization consist in dynamical neurophysiological states tonotopically distributed.

This idea is in line with fMRI, MEG, EEG and ECoG studies which showed that the auditory areas are characterized by a hierarchical layout highly specialized in analyzing different aspects of the signal: the primary auditory cortex (A1) seems engaged in the acoustic processing of the signal – where broad properties of speech sound or phonetic cues for landmarks serving to identify phoneme boundaries are computed (Stevens 2002) – while the superior temporal gyrus and the superior temporal sulcus work smoothly for encoding the acoustic patterns onto phonological features (Scott & Johnsrude 2003; Price 2012; Manca & Grimaldi 2016; Santoro et al. 2014). Thus, it seems that speech discretization is a dynamical process where transient changes from the physical spectro-temporal states to neurophysiological states generate categorial perception.

Ultimately, the discrete neural states here proposed are not static: as we have seen in Section 2, an important characteristic of oscillatory rhythms is the temporal (dynamical) way in which they are constantly nested inside other through synchronization and desynchronization processes. Such characteristic is suitable for further computational processes required to reach adequate representations of the other levels of language (lexical, syntactic, etc.), which, in turn, will be realized thanks to subsequent neural states and continuous nesting of further oscillatory bands (see Grimaldi in press). At the end, the traditional multilevel properties of language (phones, phonemes, syllables, words, syntax, etc.) hypothesized by linguistics may be seen as neural states dynamically and spatially changing in the time and resulted by online computation and representations.

In conclusion, within a neurobiological perspective the classical distinction between bottom-up processes (phonetics) reflecting acoustic differences and top-down (phonology) processes reflecting distinctive features extraction should be reinterpreted as a continuous process involving different physical states (spectro-temporal states and neurophysiological states) where a phase transition indicates a progressive changes in structure and a consequent changes in properties, so that the properties of the former states are converted in the properties of the latter states, as it happens for states of matter. With a crucial difference: changes in structure and in properties from the spectro-temporal states to the neurophysiological states generates abstract representations of speech sounds.

## Acknowledgements

I wish to thank two anonymous reviewers for their useful comments on the first version of the manuscript that have permitted me to improve the work.

